# A robust method to estimate regional polygenic correlation identifies heterogeneity in the shared heritability between complex traits

**DOI:** 10.1101/143644

**Authors:** Guillaume Paré, Shihong Mao, Wei Q. Deng

**Affiliations:** Population Health Research Institute, Hamilton Health Sciences and McMaster University, Hamilton, Canada; Population Genomics Program, Department of Clinical Epidemiology and Biostatistics, McMaster University, Hamilton, Canada; Department of Pathology and Molecular Medicine, McMaster University, Hamilton, Canada; Department of Statistical Sciences, University of Toronto, Toronto, Canada

**Keywords:** Complex traits, polygenic inheritance, genetic correlation, linkage disequilibrium, 1000 Genomes Project, maximum likelihood

## Abstract

**Background:** Complex traits can share a substantial proportion of their polygenic heritability. However, genome-wide polygenic correlations between pairs of traits can mask heterogeneity in their shared polygenic effects across loci. We propose a novel method (WML-RPC) to evaluate polygenic correlation between two complex traits in small genomic regions using summary association statistics. Our method tests for evidence that the polygenic effect at a given region affects two traits concurrently.

**Results:** We show through simulations that our method is well calibrated, powerful and more robust to misspecification of linkage disequilibrium than other methods under a polygenic model. As small genomic regions are more likely to harbour specific genetic effects, our method is ideal to identify heterogeneity in shared polygenic correlation across regions. We illustrate the usefulness of our method by addressing two questions related to cardio-metabolic traits. First, we explored how regional polygenic correlation can inform on the strong epidemiological association between HDL cholesterol and coronary artery disease (CAD), suggesting a key role for triglycerides metabolism. Second, we investigated the potential role of PPARγ activators in the prevention of CAD.

**Conclusions:** Our results provide a compelling argument that shared heritability between complex traits is highly heterogeneous across loci.

## Background

Most complex traits follow a polygenic model of inheritance, whereby thousands of common genetic variants contribute to phenotypic variance. Furthermore, genetic variance is not spread evenly throughout the genome, but rather, tends to concentrate in specific regions [1-3]. Shared polygenic heritability between pairs of complex traits has been shown at a genome-wide level, and there is broad interest in developing novel methods to estimate such shared genetic architectures between pairs of complex traits [4-6]. Existing methods for regional correlation either use individual-level data [7, 8] or test for co-localisation of single variant associations without considering polygenic inheritance [9-12]. Nonetheless, the observation that a majority of polygenic heritability lies in variants associated below genome-wide significance, coupled with the concentration of such associations at specific loci, dictates the need for a method that can estimate polygenic correlation within small (~1 Mb) regions. As each genetic region includes a different set of genes, genome-wide correlations will miss heterogeneity in the contribution of individual genes to shared heritability. A method (pHESS) [6] was recently described to estimate regional genetic correlation, but its sensitivity to misspecification of linkage disequilibrium (LD) was not explored. This is particularly important as most recent large genetic meta-analyses are trans-ethnic, such that the LD structure underlying summary association statistics is difficult to estimate. Thus, there is a need for a method to estimate regional genetic correlation that is robust to misspecification of LD.

Further motivation for this work stems from a series of observations suggesting that (1) the majority of additive genetic effects appear to be polygenic and well below genome-wide significance, (2) polygenic inheritance tends to concentrate at specific regions [1], and (3) there is widespread genome-wide correlation observed in pairs of complex traits [4]. We propose a novel method (WML-RPC) to estimate the regional polygenic correlation between two traits, retaining all variants in a given region, irrespective of LD, and using summary association statistics. Our method adopts a weighted maximum likelihood approach to estimate the regional polygenic variance of each trait and their polygenic correlation. It assumes random polygenic effects, or in other words, that multiple genetic variants are associated with a trait in each region. This framework builds on our previous work [1, 13] and has the distinct advantage of being robust to misspecification of either LD or genetic effect sizes. Unlike other approaches, our method makes no assumption about the causal relationship of one trait over the other, but rather, is intended to test whether a single polygenic effect affects two traits concurrently at a given locus. In addition, as WML-RPC provides estimates for the strength of the polygenic correlation, it can be used to test for the presence of correlation, or alternatively, for deviation from a set level of correlation (i.e., the null hypothesis can be set to any level of genetic correlation). We illustrate the utility of our method by using it to answer two questions related to cardio-metabolic traits, bringing novel insights into the inverse association of HDL cholesterol with coronary artery disease (CAD), and exploring the role of PPARγ activators in the prevention of CAD.

## Results

### Simulations using 1000 Genomes Project Haplotypes

We simulated two traits using phased 1000 Genomes (1000G) Project [14] haplotypes. The five simulated regions comprised from 296 to 1,117 SNPs, corresponding to a physical distance of ~1Mb, and summary association statistics were generated in two distinct populations of 100,000 simulated individuals each. Assuming realistic levels of genetic association, there was no type I error inflation when either or both traits were truly genetically associated in the absence of any genetic correlation at p<0.001. Based on 100,000 simulations, the proportion of false-positives was 0.00015 at a more stringent α-level of 0.0001, which did not differ significantly from the expected (p=0.083). The power to detect genetic correlation was dependent on genetic effect sizes and the strength of true underlying genetic correlation (Figure 1). We evaluated the influence of correlated error terms, which could occur if summary association statistics were derived from overlapping sets of participants. With 25% of participants overlapping and the non-genetic correlation between traits set at *ρ* = 0.2, the impact on both type I error rate and power was minimal (Supplementary Figures S1 and S2) at p<0.001. We also tested a more extreme scenario that assumed a complete overlap in participants assuming a correlation of non-genetic error terms of 0.2. Again, minimal type I error inflation was observed under the null hypothesis of no correlation (Supplementary Figures S1 and S2) at p<0.001.

**Figure 1:**
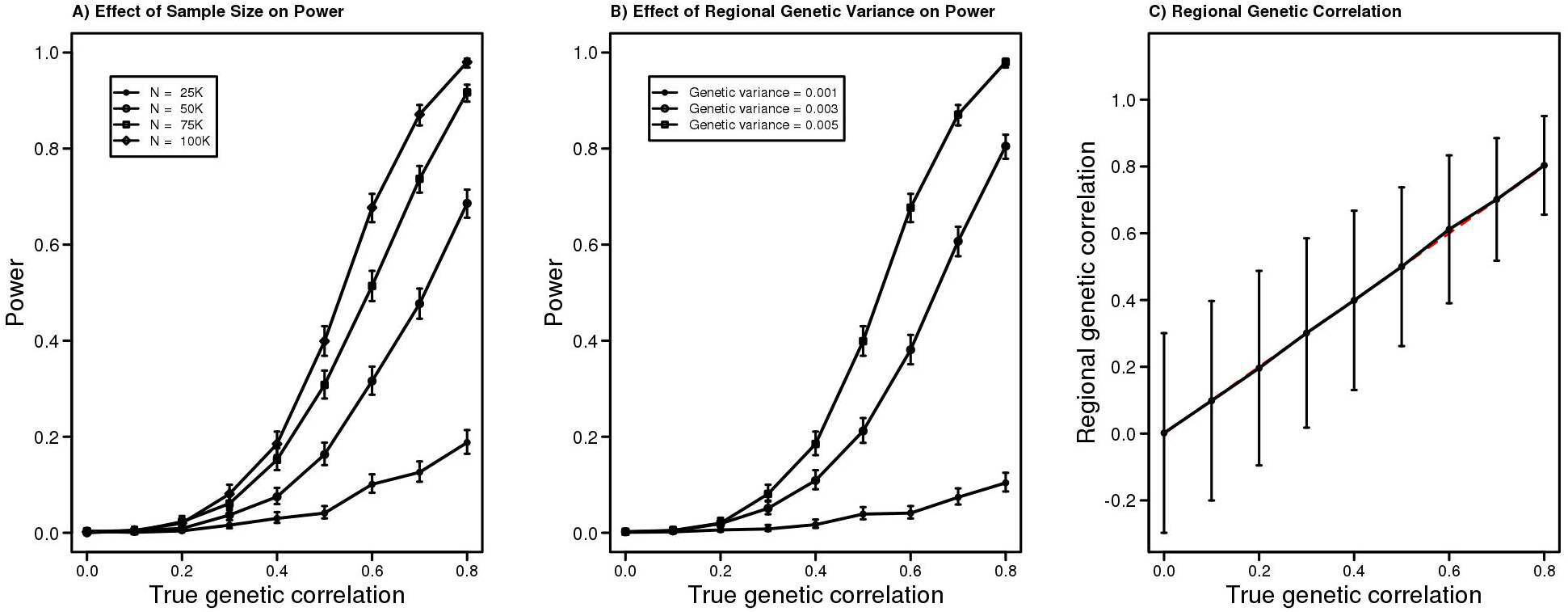
Performance of WML-RPC in simulated data using 1000 Genomes Project haplotypes. The power to detect polygenic correlation at a nominal α-level of 0.001 as a function of the true polygenic correlation was calculated over 1,000 simulated replicates on a region of 1Mb simulated using haplotypes of European participants from 1000 Genomes Project. In panel A), the sample size ranged from 25K to 100K individuals, while keeping the true regional genetic variance constant at 0.005 for each trait. In panel B), sample size was fixed at 100K individuals, but the regional polygenic variance varied from 0.001 to 0.005. In panel C), the mean regional polygenic correlation over 1,000 replicates is illustrated as a function of the true (red dashed line) polygenic correlation, assuming a sample size of 100K and genetic variance of 0.005 for both traits. The error bars represent the mean polygenic correlation ± 1.96SD over 1,000 replicates.

We also sought to benchmark our method against other recently described co-localisation methods. Two of the tested methods (gwas-pw [12] and jlim [10]) assess the possibility that a single causal variant underlies a genetic association with two traits. As expected given their model assumptions, neither method performed well in the presence of polygenic inheritance, with both showing inflated type I error rates and decreased power compared to WML-RPC (Figure 2). On the other hand, pHESS [6] was designed to assess genetic correlation under a polygenic model and its power was better than that of WML-RPC (Figure 2A), although at the expense of inflated type I error rates (0.011 and 0.0020 at p-value thresholds of 0.001 and 0.0001, respectively) under the null hypothesis of no correlation. To assess the robustness of WML-RPC and pHESS to misspecification of LD structure, additional analyses were performed, as previously, but with the reference LD matrix calculated using an increasing proportion of 1000G African (AFR) individuals (Figure 2). In other words, summary association statistics were calculated using only individuals of European ancestry under the null hypothesis of no genetic correlation while the LD matrix used for WML-RPC and pHESS calculations included varying proportions of African individuals. As shown in Figure 2B, WML-RPC type I error was minimally affected by differences in LD structure, even under scenarios of extreme LD misspecification. However, pHESS was more sensitive to LD misspecification with type I error increasing up to 0.161 using a p-value threshold of 0.001 (Figure 2B).

**Figure 2:**
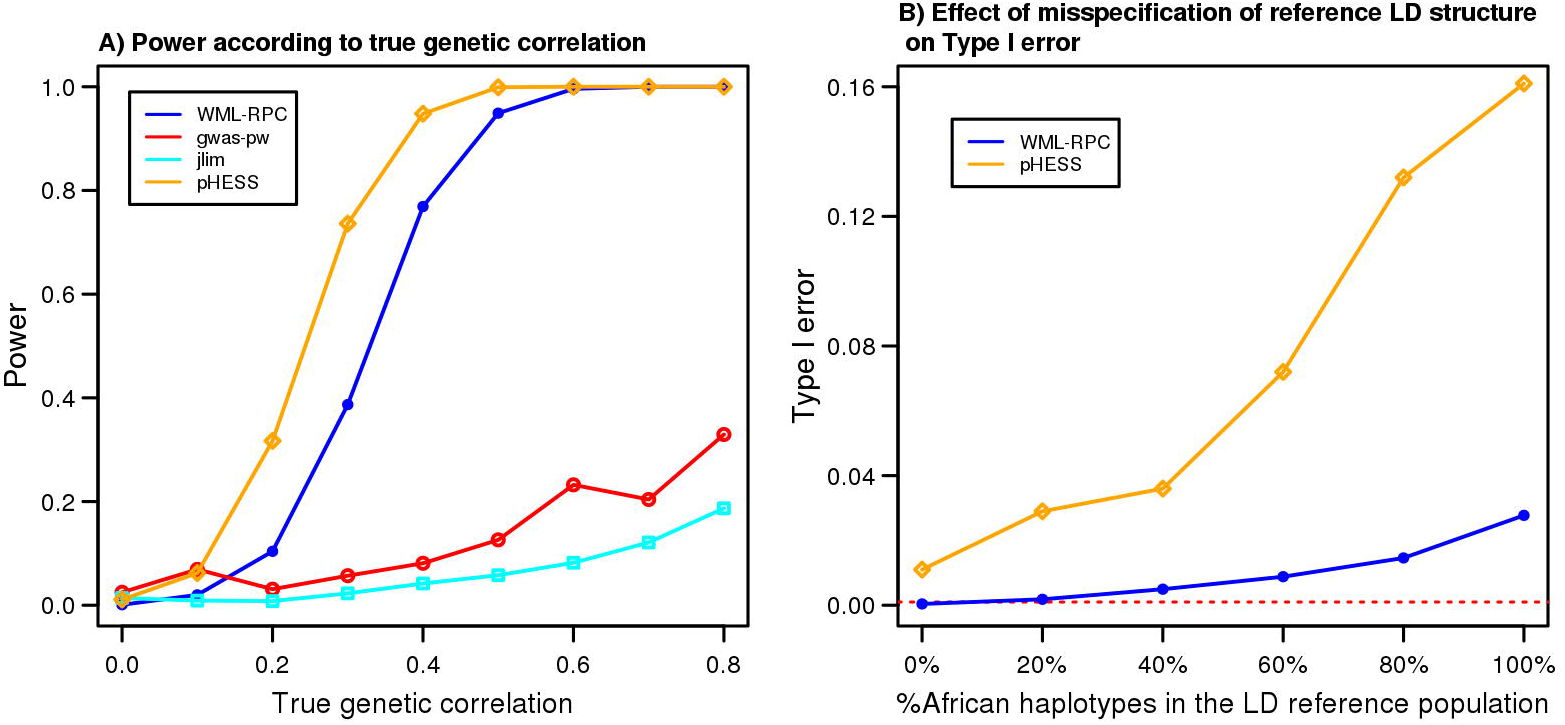
A comparison of statistical power to detect true genetic correlation and type I error resilience by WML-RPC, gwas-pw, jlim, and pHESS. A) Power to detect polygenic correlation at a nominal α-level of 0.001 (for WML-RPC, jlim, and pHESS) or posterior probability > 0.90 (for gwas-pw) as a function of the true polygenic correlation. Results are based on regions of 1Mb simulated using haplotypes of European participants from 1000 Genomes Project, with a sample size of 100,000 and assuming the true regional genetic variance is 0.005 for each trait. For each condition, the simulation was repeated 1,000 times unless stated otherwise. Under the null hypothesis of no polygenic correlation, the type I error rate, which can be assessed at the true genetic correlation of 0, was 0.1% for WML-RPC (10,000 simulations), while the estimated rates of type I error were 2.5%, 1.4%, and 1.1% for gwas-pw, jlim, and pHESS, respectively. B) Comparison of the effect of misspecification of the reference LD structure on type I error between WML-RPC and pHESS methods. Results are based on simulations performed under the same parameters described in A); however, the LD matrix was calculated using an increasing number of individuals of African ancestry. The dashed red line represents the expected type I error at a nominal α-level of 0.001.

WML-RPC and pHESS were also used to calculate the regional genetic correlation under different LD structures. As shown in Figure 3, regional genetic correlation estimates from both methods agreed with the true genetic correlation, even when a gross misspecification of LD structure was applied (Figure 3B). However, the dispersion of estimates was larger with pHESS than WML-RPC when LD was misspecified, as illustrated by larger confidence intervals.

**Figure 3:**
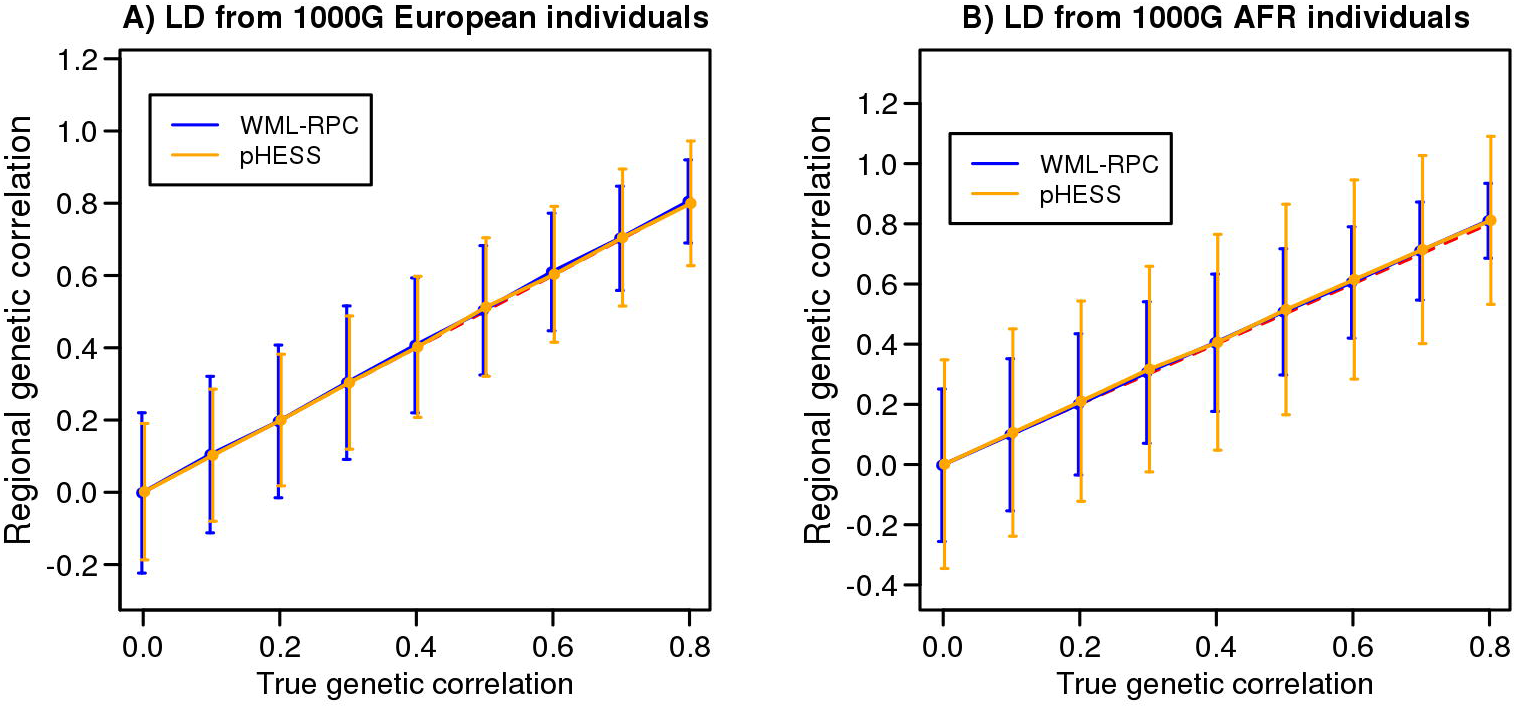
Comparison of the estimated regional polygenic correlation by WML-RPC and pHESS methods under different LD structures. Results are based on regions of 1Mb simulated using haplotypes of 1000 Genomes Project participants of European descent, with a sample size of 100,000 and assuming the true regional genetic variance is 0.005 for each trait and non-genetic correlation of 0. Regional genetic correlations were calculated under the LD structure for A) European (i.e., the correct LD specification) and B) African individuals (i.e., a gross misspecification of LD).

### Insights into the relationship between HDL cholesterol and coronary artery disease

To illustrate how regional polygenic correlation can provide novel epidemiological insights, we first explored the genetic relationship between HDL cholesterol (HDLc) and CAD using summary association statistics from large genetic meta-analyses. Blood HDLc concentration is one of the strongest predictors of decreased CAD risk in epidemiological studies [15], yet it is widely agreed that the association is non-causal. Several Mendelian randomization studies have been conducted to address this question, supporting a lack of causal relationship [16-20]. Furthermore, pharmacological interventions to raise HDLc have thus far been disappointing [21-23], further strengthening the hypothesis of a non-causal relationship. If the relationship is truly non-causal, then one or more upstream biological pathways can be expected to jointly affect HDLc concentration and the risk of CAD; thus, explaining the strong epidemiological association. In other words, underlying causal risk factor(s) must exist that cause decreased HDLc to increase the risk of CAD, even if HDLc itself is an epiphenomenon. Regional polygenic correlation can help identify regions whose effects on HDLc and CAD are consistent with epidemiological studies, and thus, provide insights into the identity of biological pathways that are responsible for their strong epidemiological association.

We divided the genome into 2,687 regions of ~ 1Mb and determined which regions showed evidence of polygenic correlation between HDLc and CAD. Keeping only the 848 regions with at least nominal evidence (p < 0.05) of a polygenic association with either HDLc or CAD, we tested for regional polygenic correlation and applied a conservative Bonferroni correction (p <0.05/2,687). Consistent with a non-causal role of HDLc in CAD, none of the seven regions identified were directly involved in HDL production (e.g., the APOA1 locus) and heterogeneity in polygenic correlation was present, with one region having positive polygenic correlation while others had negative correlation (Table 1 and Supplementary Table S1 for unclipped 95% confidence intervals for regional polygenic correlations). The seven regions with significant negative polygenic correlation between HDLc and CAD are of particular interest since they could potentially underpin the epidemiological association. Tellingly, all of the seven regions were located at loci directly related to triglycerides metabolism. The *LPL, TRIB1* and *MC4R* loci are strongly associated with fasting triglycerides [24], while *APOE* is linked to the postprandial regulation of triglyceride-rich lipoproteins [25] and SORT1 to hepatic triglyceride-rich VLDL secretion [26]. The region with significant (*p* = 1.8 x10^−6^) positive correlation encompassed the gene encoding hepatic lipase (*LIPC*). *LIPC* deficiency leads to increased HDLc [27] and triglycerides-rich intermediate-density lipoproteins (IDL) [28], but its role in CAD remains controversial. Consistently, genome-wide significant positive polygenic correlation between HDLc and triglycerides was also observed at the locus surrounding *LIPC* (data not shown).

**Table 1:** Regions with significant polygenic correlation estimates between HDLc and CAD.

| CHR | Position (Kb) | Candidate gene | Candidate Gene function | HDLc polygenic regional *p*-value | CAD polygenic regional *p*-value | Polygenic correlation (95% CI) | Polygenic correlation *p*-value |
| --- | --- | --- | --- | --- | --- | --- | --- |
| 1 | 109,723-110,710 | SORT1 | VLDL secretion | 2.58E-24 | 3.16E-20 | -0.83(-1.00, -0.61) | 3.69E-07 |
| 8 | 18,869-19,875 | LPL | Lipoprotein triglyceride lipase | <1.00E-323 | 2.09E-06 | -0.94 (-1.00, -0.79) | 1.02E-10 |
| 8 | 19,875-20,875 | LPL | Lipoprotein triglyceride lipase | 1.86E-213 | 5.81E-04 | -0.93(-1.00, -0.79) | 5.17E-08 |
| 8 | 126,042-127,062 | TRIB1 | Regulation of hepatic lipogenesis | 1.43E-48 | 1.66E-03 | -1.00 (-1.00, -0.79) | 2.84E-06 |
| 15 | 58,311-59,348 | LIPC | Hepatic triglyceride lipase | <1.00E-323 | 9.49E-02 | 1.00 (0.87, 1.00) | 1.82E-06 |
| 18 | 57,456-58,421 | MC4R | Appetite regulation | 5.61E-18 | 1.12E-06 | -0.98(-1.00, -0.79) | 1.07E-05 |
| 19 | 44,789-45,840 | APOE | Catabolism of triglyceride-rich lipoproteins | 6.83E-52 | 2.62E-10 | -0.89 (-1.00, -0.77) | 6.74E-11 |

The genome-wide genetic correlation between HDLc and CAD was estimated at −0.44 when using a weighted average of regional genetic correlations, while it was estimated at −0.25 (0.07) with LDScore regression [4]. The distribution of regional genetic correlations is illustrated in Supplementary Figure S3.

Overall, our results support the hypothesis the role of HDLc, as a marker of triglycerides levels, can help explain the strong epidemiological association with CAD. Triglycerides are known causal mediators of CAD [20]. Their levels are notoriously variable and can increase dramatically in the post-prandial state. As HDLc concentrations are more stable and inversely correlated to triglycerides concentrations, they can provide a surrogate for long-term exposure to triglycerides. Indeed, non-fasting triglycerides, although seldom measured, have been shown to better predict CAD risk than fasting measurements [29]. Our results also suggest that high HDLc caused by decreased *LIPC* activity increases the risk of CAD.

### Thiazolidinediones, PPARγ and risk of CAD

Pharmacological activation of PPARγ with thiazolidinediones is used to treat and prevent diabetes. However, the role of thiazolidinediones in the prevention of CAD is controversial. *Post hoc* analyses of randomized trials identified the potential for thiazolidinediones to increase CAD risk [30], which, with the exception of pioglitazone, led to the removal of all thiazolidinediones from clinical use. Based on these observations, a large clinical trial addressing the issue of CAD risk reduction by rosiglitazone was stopped early [31]; thus, leaving this important clinical question unanswered. The controversy was further fuelled by the recent publication of the IRIS trial showing a significant *reduction* in cardiovascular events in individuals randomized to pioglitazone [32]. While the ongoing debate around the cardiovascular protective effects of thiazolidinediones has focused on their glucose-lowering effects, our results suggest their effect on lipoproteins (also seen in randomized clinical trials) might be of greater importance with respect to CAD than previously appreciated. We tested the region surrounding *PPARG* (+/- 500 Kb) for evidence of association with cardiometabolic traits. As expected from the known pharmacological effects of thiazolidinediones [33], significant (*p* < 0.05) regional associations were observed with diabetes, triglycerides, HDLc, LDLc and BMI. As an added measure of sensitivity, assessments of +/- 100 Kb and +/- 300 Kb regions surrounding the *PPARG* locus were also performed. The reduction in assessment region yielded similar results to those observed when a +/- 500 Kb region was used (Supplementary Tables S2 and S3, respectively)

We then tested this set of traits for polygenic correlation with diabetes and CAD (Table 2). The polygenic variance p-values at the *PPARG* locus are shown in Table 2 (column 2). A significant (p<0.05/9) and positive polygenic correlation was observed between diabetes and triglycerides, LDLc and CAD, and triglycerides and CAD. A trend towards a negative polygenic correlation was observed between diabetes and BMI (p=0.008). Polygenic correlation was not significant between diabetes and LDLc, or diabetes and CAD. Polygenic correlation between LDLc and CAD is of particular interest since pioglitazone has recently been shown to reduce LDL particle number and size [34, 35]. This observation and the polygenic correlation with triglycerides support the hypothesis that the protective effect of pioglitazone (and perhaps other thiazolidinediones) on CAD risk is the consequence of its beneficial effect on atherogenic lipoproteins.

**Table 2:**
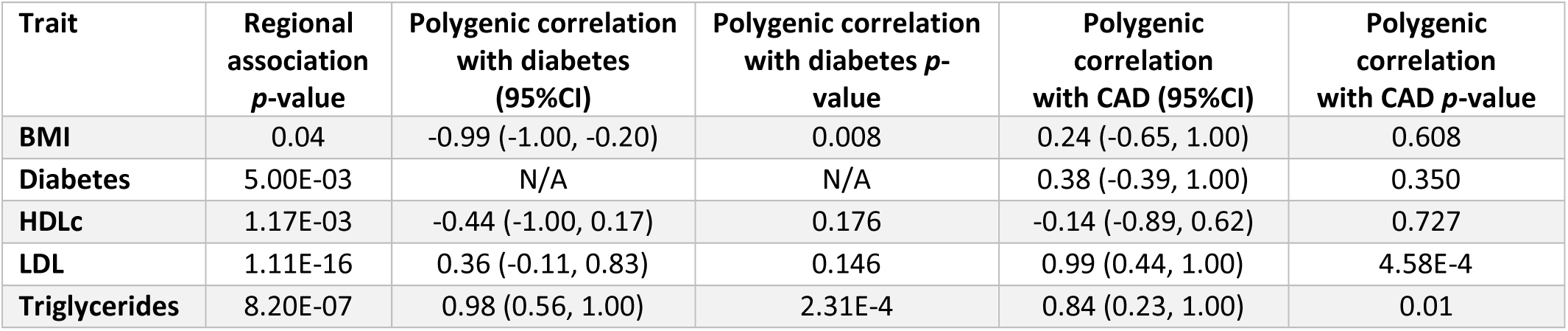
Polygenic correlation at the *PPARG* locus.

## Discussion

We herein propose a novel method to estimate regional polygenic correlation between two traits. Our method is distinct from other co-localisation and genetic correlation tests as it is based on a polygenic model of inheritance, only requires summary association statistics, and is robust to misspecification of the LD structure. The latter point is of particular importance as large genetic meta-analyses include participants of mixed ancestry, such that the underlying LD structure is difficult to estimate. Our approach is particularly attractive when studying complex traits with strong polygenic inheritance, where any single genetic association is unlikely to fully capture a large proportion of genetic effects. Our method has several other advantages, including the ability to adjust for LD, the possibility to test specific hypotheses regarding polygenic correlation, and the ability to estimate regional genetic variance for a single trait. WML-RPC has wide ranging applications, as we have illustrated. It can help discover biological pathways explaining epidemiological associations such as those for HDLc and CAD, identify regions with complex patterns of polygenic correlation, or help gain insights into the role of single genes or drug targets.

Our examples make a compelling argument that shared heritability is highly dependent on regional genetic effects. Unless a locus has a direct effect on a risk factor (e.g., the APOB or LDLR loci on LDLc), it cannot be assumed that correlation implies a causal effect of the risk factor on the outcome. For instance, genetic correlation between HDLc and CAD at the *LIPC* locus, combined with prior knowledge of the effect of *LIPC* on intermediate density lipoproteins, suggests that decreased *LIPC* activity leads to both increased HDLc and CAD risk. Inclusion of that locus in Mendelian randomization studies may thus result in biased inferences about the causal role of HDLc in CAD. Such considerations stress the importance of taking the biological effects of each genetic region into account before concluding on the relationship between a risk factor and outcome. Knowledge of biological effects can also provide insights into epidemiological relationships, such as regions with negative correlation between HDLc and CAD pointing to triglycerides metabolism as a key factor to explain the epidemiological association.

WML-RPC can also be used to explore candidate gene regions. We found that regional polygenic associations recapitulate the effects of PPARγ agonist thiazolidinediones on cardio-metabolic traits. Our results support the hypothesis thiazolidinediones can reduce CAD risk through their effect on lipids, particularly LDLc and triglycerides. In line with this hypothesis, recent data have shown that pioglitazone decreased the concentration of atherogenic lipoproteins [34, 35]. However, genetic correlation with CAD was only significant with LDLc and triglycerides, but not diabetes itself, as might have been expected given triglycerides had significant correlation with diabetes and CAD. While this finding could have been serendipitous, it is also possible that genetic variants regulating *PPARG* function vary from one tissue to the other, such that genetic regulation of LDLc at the *PPARG* locus (and thus, the risk of CAD) overlaps only partially with its effect on diabetes. Indeed, such tissue-specific effects of *PPARG* have been described [33], with adipocytes being mainly responsible for glycemic effects and hepatocytes regulating atherogenic lipoprotein metabolism [36]. Similarly, it is possible that thiazolidinediones have varying affinities for different tissues. This illustrates a further advantage of our method as it is agnostic to gene regulation mechanisms, and thus, not dependent on known eQTL associations, which may vary according to tissue and cellular context.

There are some limitations to this study. First, the WML-RPC approach assumes that all functional variants within a region affect a trait through the same pathway. Importantly, distinguishing a single causal variant for multiple traits, as opposed to multiple causal variants, in the presence of strong LD, is not possible. Thus, the presence of two highly correlated causal variants is also possible and should be considered [12]. Second, some loci may fail to fit a polygenic model, for example when there is a single very strong association at a locus, and other methods might be better suited. Third, statistical power to detect genetic correlation depends on sample size and genetic variance. While we are confident in regions identified using stringent statistical criteria, many other covariant regions have likely been missed. Fourth, many regions have no known candidate genes, in which case our method can point to regions of interest, but not necessarily biological interpretations. Nonetheless, improving knowledge of gene function and regulation, combined with the expanding repertoire of genome-wide association studies, should provide increasing opportunities for WML-RPC to lead to novel insights into complex traits. Fifth, reliable correlation estimates depend on the quality of summary association statistics, for instance, the proper adjustment for population stratification in large genomic consortia, which is almost invariably the case.

## Conclusions

In conclusion, we present a novel robust method to estimate regional polygenic correlation using summary association statistics. WML-RPC can estimate polygenic correlation within relatively small genetic regions, enabling a more detailed characterization of genetic correlation than genome-wide genetic correlations. Our method can be used to identify pathways shared between two traits, pinpoint regions of interest, or test specific hypotheses for a given gene. Our examples illustrate the heterogeneity in pairwise genetic correlation across loci. They support the notion that genetic effects are specific to each region and that unless a locus directly affects a risk factor, caution must be exercised when making causal inferences.

## Methods

### Overview of Methods

Here, we propose a novel weighted maximum likelihood (WML) method to estimate the regional polygenic correlation between pairs of complex traits using summary association statistics. Our framework is derived from our previous work [13] where we developed a simple procedure to estimate regional polygenic variance for single traits using summary association statistics. We assume random, normally distributed genetic effects for each trait. The polygenic regional effects for each trait under the random effects model was first estimated, and then the polygenic correlation between traits was estimated using WML. Our method has several advantages, including a convenient framework for hypothesis testing using a likelihood ratio test, the use of summary-level data, and robustness to misspecification of LD structure. We validated our method using data simulated based on the 1000 Genomes Project and applied our method to summary-level data from large genetic meta-analyses of cardio-metabolic traits.

### Estimation of regional polygenic variance and correlation

We recently described a simple procedure to estimate regional genetic variance using summary association statistics, adjusting for linkage disequilibrium (LD) [13]. We now propose to adapt this procedure to estimate polygenic correlation between a pair of traits using a weighted maximum likelihood (WML) approach. Suppose the genotype matrix is fixed while the true, unobserved genetic effect is a random vector β, whose individual components, i.e., the effect size of SNPs, *I* = *1,2,…m*, have mean zero and variance *σ*^2^. The true, unobserved, genetic model can be expressed as:

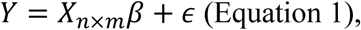

where *ε* is a vector of standard normal error with identity variance-covariance matrix and the genetic variance is given by *mσ*^2^. Without loss of generalizability, we assume the observed quantitative trait (*y*) and the *n* x *m* genotype matrix *X* standardized to have zero mean and unit variance throughout. The pairwise LD (*r^2^*) between two SNPs *k* and *l* is denoted by 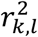. For a SNP *d* the following LD adjustment (*η*_*d*_) can be defined as the summation of LD between the *d*^th^ SNP and all SNPs within the same region (herein defined as encompassing all SNPs within ~1 Mb):

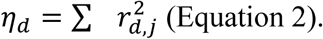

We suggest setting all instances of 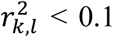 to zero when considering large regions to avoid imprecision in the estimation of 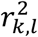 unduly inflating *η*_*d*_. Only including SNPs with summary GWAS statistics in the sum, the estimated variance explained by each SNP *d* is given by:

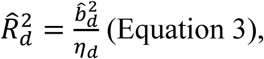

where 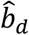 denotes the univariate regression coefficient commonly reported in GWAS results with sample size *N* (assuming genotypes from external GWASs have also been standardized to have zero mean and unit variance). Assuming a strictly additive genetic model where each SNP contributes additively to a trait without any interaction or haplotype effects, we have previously shown[13] that 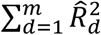 is an estimator of the regional variance *mσ*^2^ by demonstrating the approximated equivalence between the expected total genetic variance over a region

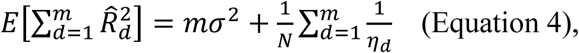

and the estimated adjusted coefficient of determination in multiple linear regression can be considered an estimator of the regional variance

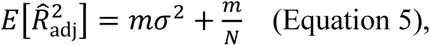

when the sample size is sufficiently large.

Since the true genetic effects are given by a random vector *β*, this implies:

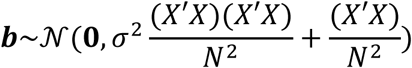

or marginally:

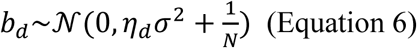

As we are interested in the estimation of *σ*^2^ via the surrogate 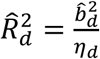, the following weighted likelihood function is maximized to find 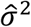:

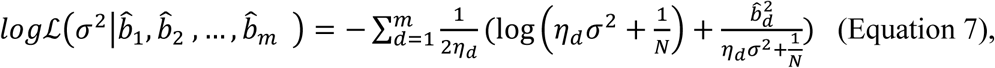

where the log-likelihood of each observed 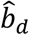 is weighted by the inverse of the LD adjustment, such that if two SNPs were in complete LD, then effectively only one SNP contributes to the log-likelihood for the genetic variance.

This framework can be extended to study the genetic correlation between a pair of traits.

In this scenario we have 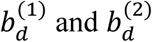, which are the summary association statistics for trait 1 and 2, respectively, following a bivariate normal distribution:

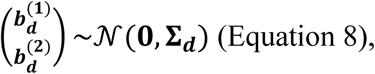

with covariance matrix:

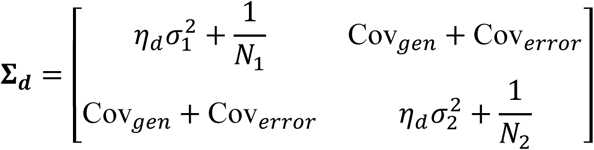

where 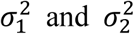 are the genetic variance of trait 1 and 2, respectively, and *N*_1_ and *N*_2_ the corresponding sample sizes. Cov_*gen*_ represents the genetic covariance between both traits whereas Cov_*error*_ is the error term covariance and can be assumed to be zero.

The weighted likelihood function can be adapted using genetic variance estimates from the previous weighted likelihood 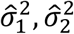:

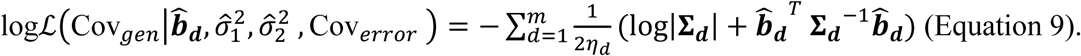

The maximum likelihood estimates of Cov_*gen*_ enables the use of a likelihood ratio test for hypothesis testing. While Cov_*error*_ could be estimated, we found that under realistic scenarios its effect is negligible and has therefore been set to zero for current analyses. This might not be ideal when the correlation of error terms is very strong, in which case a non-zero Cov_*error*_ could be used in the likelihood estimation. As a note, Cov_*gen*_ estimated from empirical data can cause numerical estimates to be higher than both 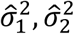 or lower than 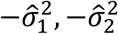 and thus causing the genetic correlation Cor_gen_ to be higher 1 or lower than −1, and will correspondingly be set to 1 or −1. Also note that stable and meaningful estimates of Cor_gen_ can only be obtained when both traits 1 and 2 have positive estimates for regional genetic variance. We also note that the stability of genetic correlation estimates depends on the regional genetic effect size of both traits, such that stable estimates can be obtained when one trait is weakly associated while the other trait is strongly associated. Finally, although the aim of the method is to estimate regional polygenic correlation, it is important to confirm there is no bias in regional polygenic variance under the null hypothesis of no genetic association, which we checked using simulations (Supplementary Figure S4).

### Simulations using 1000 Genomes Project data and a study of cardio-metabolic traits

We used 379 participants of European descent from the 1000G [14] as the reference panel for LD as it is the dominant ancestry in the studies included. Phased haplotypes are provided by the 1000G project for each of the 379 participants (i.e., a total of 758 phased haplotypes). The set of 758 haplotypes constitutes the reference population, with each haplotype having an equal allele frequency of 1/758=0.0013. We randomly sampled two phased haplotypes (with replacement) from the 758 phased haplotypes for each of the 100,000 simulated individual, and derived corresponding genotypes. For each simulated individual, two traits were simulated based on unobserved genetic effects and error terms. The simulated traits were used to derive the summary association statistics for each SNP by regressing the simulated traits on SNP genotype (i.e., summary association statistics were simulated, not fixed). Since polygenic inheritance was being assessed, we used an infinitesimal model according to which every SNP was associated to some extent, and the effect size was sampled from a normal distribution. Thus, the set of causal SNPs was not the same for both traits, with the exception of scenarios where both traits had perfect regional genetic correlations. The LD structure needed as input for calculation of regional genetic correlation was derived from an independent set of 1,000 simulated individuals, again using the 758 phased European haplotypes as the reference population. To assess the robustness of methods to misspecification of LD structure, we included a varying proportion of 1000G phased haplotypes of African descent in the reference population when simulating the 1,000 individuals. All simulations were performed on five randomly chosen regions of 1 Mb. Only the results from the first region are presented in the main manuscript, but consistent results were obtained for all four other regions (Supplementary Figure S5).

We tested our method using summary association statistics from large genetic meta-analyses of cardiometabolic traits, including coronary artery disease [37], LDL cholesterol, HDL cholesterol, triglycerides [38], type 2 diabetes [39], body mass index [40], and blood pressure [41]. We identified a common set of SNPs among all corresponding meta-analyses and subsequently divided the genome into blocks of ~1Mb by calibrating the number of SNPs per block to minimize inter-block LD, as previously described [13].

## List of abbreviations

1000G – 1000 Genomes Project

AFR – African

APOA1 – Apolipoprotein A1

APOE – Apolipoprotein E

BMI – Body-mass index

CAD – Coronary artery disease

GWAS – Genome-wide association studies

HDL – High-density lipoprotein

HDLc – HDL cholesterol

IDL – Intermediate-density lipoproteins

Kb - Kilobases

LD – Linkage disequilibrium

LDLc – Low-density lipoprotein cholesterol

LIPC – hepatic lipase C

LPL – Lipoprotein lipase

Mb - Megabases

PPARG – Peroxisome proliferator-activated receptor gamma

SNP – Single nucleotide polymorphism

TRIB1 – Tribbles pseudokinase 1

TG - triglyceride

WML – Weighted maximum likelihood

## Declarations

Availability of data and materials

The datasets generated and/or analysed during the current study are available from the corresponding author on reasonable request.

## Competing interests

The authors declare that they have no competing interests.

## Funding

Funding for this study was provided by the Canada Research Chair program as well as the Cisco Professorship in Integrated Health Systems.

## Authors’ contributions

GP conceived and designed the study, analysed and interpreted the data, and wrote the manuscript. SM developed the analytical software, performed analyses, analysed the data and wrote the manuscript. WQD performed analyses, analysed and interpreted the data, and wrote the manuscript. All authors read and approved the final manuscript.

---

Table S1: Regions with significant polygenic correlation estimates (unclipped) between HDLc and CAD.

Table S2: Regional polygenic correlation at the *PPARG* locus (+/- 100Kb).

Table S3: Regional polygenic correlation at the *PPARG* locus (+/- 300Kb).

**Figure S1: Quantile-quantile plot of WML-RPC correlation *p*,-values against the null hypothesis of no polygenic correlation.**

Quantile-quantile plot of polygenic correlations-values from WML-RPC on a region of 1Mb with 100K individuals simulated using haplotypes of European participants from the 1000 Genomes Project. Each condition was repeated 1,000 times. The observed-log10 *p*-values were plotted against the expected, shown with 95% confidence regions (dashed red lines). We assumed regional genetic variances of 0.005 for each trait, without polygenic correlation. In panel A), we assumed the error terms were uncorrelated with *ρ* = 0. In panel B), 25% of participants overlapped between traits, such that the non-genetic correlation was set at *ρ* = 0.05. In panel C), the overlap between participants was set at 100%, again with the non-genetic correlation set at *ρ* = 0.2.

**Figure S2: Statistical power and estimated polygenic correlation as functions of the true polygenic correlation in simulated data when error terms correlate.**

The power of WML-RPC to detect polygenic correlation at a nominal α-level of 0.001 for each true polygenic correlation values was calculated over 1,000 simulated replicates on a region of 1Mb simulated using haplotypes of European participants from the 1000 Genomes Project. The non-genetic correlation between error terms was set at 0.05 in simulations illustrated in panels A), B) and C), and was set to 0.2 in simulations illustrated in panels D), E) and F). In panels A) and D), the sample sizes varied from 25K to 100K individuals, while keeping the true regional genetic variance constant at 0.005 for each trait. In panels B) and E), the sample size was constant at 100K individuals, but the true regional genetic variance was varied from 0.001 to 0.005. In panels C) and F), the mean (±SD) estimated regional polygenic correlation is illustrated in black, as a function of the true (red dashed line) polygenic correlation, assuming a sample size of 100K and a true genetic variance of 0.005 for both traits.

**Figure S3: Distribution of estimated regional genetic correlation between HDLc and CAD.**

Regional genetic correlations for 2,687 regions were estimated for high-density lipoprotein cholesterol (HDLc) and coronary artery disease (CAD).

**Figure S4: Quantile-quantile plot for regional genetic variance *p*-values generated by WML-RPC under the null hypothesis of no regional genetic variance.**

Each observed p-value was calculated based on a region of 1Mb simulated using haplotypes of 1000 Genomes Project participants of European descent, with a sample size of 100K and assuming the true regional genetic variance was 0. The simulation was repeated 1,000 times. The observed *p*-values were plotted on a-log10 scale against the expected, shown with 95% confidence regions in dashed red lines.

**Figure S5: Performance of WML-RPC in simulated data using 1000 Genomes Project haplotypes at five other randomly chosen regions.**

The power to detect polygenic correlation at a nominal α-level of 0.001 as a function of the true polygenic correlation was calculated over 1,000 simulated replicates on five regions of 1Mb each, simulated using haplotypes of European participants from 1000 Genomes Project. In panels A), C), E), G), I) the sample sizes ranged from 25K to 100K individuals, while keeping the true regional genetic variance constant at 0.005 for each trait. In panels B), D), F) H), J), the mean regional polygenic correlation over 1,000 replicates is illustrated as a function of the true (red dashed line) polygenic correlation, assuming a sample size of 100K and genetic variance of 0.005 for both traits. The error bars represent the mean polygenic correlation ± 1.96SD over 1,000 replicates.

